# The extracellular sialidase NEU3 induces neutrophil priming

**DOI:** 10.1101/2022.02.23.481673

**Authors:** Sara A. Kirolos, Richard H. Gomer

## Abstract

Some extracellular glycoconjugates have sialic acid as the terminal sugar, and sialidases are enzymes that remove this sugar. Mammals have four sialidases, but their biological functions are unclear. In this report, we show that incubation of human neutrophils with the human sialidase NEU3, but not NEU1, NEU2 or NEU4, inducess human male and female neutrophils to change from a round to a more amoeboid morphology, causes the primed neutrophil markers CD66, CD11B, and CD18 to localize to the cell cortex, and decreases the localization of the unprimed neutrophil markers CD43 and CD62L at the cell cortex. NEU3, but not the other 3 sialidases, also causes human male and female neutrophils to increase their F-actin content. The inhibition of NEU3 by the NEU3 inhibitor 2-acetylpyridine attenuated the NEU3 effect on neutrophil morphology, indicating that the effect of NEU3 is dependent on its enzymatic activity. Together, these results indicate that NEU3 can prime human male and female neutrophils, and that NEU3 is a potential regulator of inflammation.

## Introduction

An immune response is an essential process that allows the host organism to defend against pathogens (Chen, Liu, et al. 2018; Medzhitov 2007; Cekici et al. 2014). Inflammation is a tissue pathological process for the purpose of resolving infection and spurring tissue repair (Chen, Deng, et al. 2018; Walsh and Pearson 2001). Numerous immune cells reside in tissues where the cells contribute to immune defense and tissue homeostasis (Sun et al. 2020). An immune response is comprised of two distinct categories, innate and adaptive (Smith, Rise, and Christian 2019; Carsetti et al. 2020). The innate immune response is the first line of defense against foreign invaders (Kumar, Kawai, and Akira 2009; Koyama et al. 2008) and involves various cell types, including macrophages, monocytes, and neutrophils (Joseph et al. 2004; Chiu and Bharat 2016; Kobayashi et al. 2005).

In the lungs, neutrophils are a critical component of the innate immune response against pathogens during acute inflammation (Tull et al. 2009; Ding et al. 2021). Circulating neutrophils are quiescent and unprimed (Mayadas, Cullere, and Lowell 2014). As neutrophils are exposed to local activating agents, such as N-formylmethionyl-leucyl-phenylalanine (fMLF), tumor necrosis factor alpha (TNF-α), and various chemokines (Videm and Strand 2004; Hidalgo et al. 2015; Reumaux et al. 2006), priming is initiated. The priming of neutrophils is characterized by neutrophil expression of adherence molecules which allow them to attach to the endothelial cell lining of blood vessels (Tull et al. 2009). The attachment of primed neutrophils to endothelial cells ensures maximum activation of neutrophils in a highly regulated manner (Miralda, Uriarte, and McLeish 2017; Bass et al. 1986). When neutrophils bind to the endothelial cell lining, the cells begin to undergo rolling (Lawrence and Springer 1991). Although rolling does not require the activation of neutrophils, neutrophil trans-endothelial migration involves the neutrophils developing amoeboid locomotion to move into the tissue, and this is dependent on the neutrophil exposure to activating stimuli (Lawrence and Springer 1991; Diacovo et al. 1996).

As neutrophils enter a site of inflammation, neutrophils begin to release neutrophil extracellular traps (NETs), cytokines, and exosomes, and then undergo autophagy, resulting in fibroblast activation and formation of the extracellular matrix (Perez-Figueroa et al. 2021; Kolonics et al. 2021; Scapini and Cassatella 2014; Nauseef and Borregaard 2014; Ding et al. 2021). Many pulmonary disorders, such as asthma, chronic obstructive pulmonary disease (COPD), cystic fibrosis (CF), and idiopathic pulmonary fibrosis (IPF), are characterized by an influx of neutrophils into the lung tissue (Ding et al. 2021; Richter et al. 2011; Nathan et al. 2020; Barnes 2016; Yang et al. 2017), although for IPF the role of neutrophils is unclear (Ishikawa, Liu, and Herzog 2021).

Sialidases are enzymes that remove terminal sialic acids from glycoconjugates (Smutova et al. 2014; Pshezhetsky and Ashmarina 2013; Miyagi et al. 2012), and sialidases are upregulated in IPF (Karhadkar et al. 2017; Lambre et al. 1988).There are four isoforms of sialidases in mammals, NEU1, NEU2, NEU3 and NEU4 (Pshezhetsky and Ashmarina 2013), and of these, NEU3 is the only one that is primarily extracellular (Zanchetti et al. 2007). Upregulation of NEU3 is associated with multiple disorders including intestinal inflammation and colitis (Yang et al. 2021), neuroinflammation (Demir et al. 2020), periodontal disease (Shah, Thomas, and Mehta 2017) and lung fibrosis (Karhadkar, Meek, and Gomer 2021; Karhadkar et al. 2017). NEU1, NEU2 and NEU3 are upregulated in fibrotic lesions in human and mouse lungs (Karhadkar, Chen, and Gomer 2020; Karhadkar et al. 2017; Luzina et al. 2016), but only NEU3 was detected in the lung fluid of mice with bleomycin-induced pulmonary fibrosis (Karhadkar et al. 2017). Mice lacking NEU3 (*Neu3*^*-/-*^) had attenuated inflammation and fibrosis after bleomycin compared to wildtype mice (Karhadkar, Chen, and Gomer 2020). The NEU3 inhibitors 2,3-didehydro-2-deoxy-*N*-acetyl-neuraminic acid (DANA), Oseltamivir phosphate (Tamiflu), 2-acetyl pyridine (2AP), methyl picolinate (MP), and 4-Amino-1-methyl-2-piperidinecarboxylic acid (AMPCA) also reduced bleomycin-induced lung fibrosis in mice (Karhadkar, Meek, and Gomer 2021; Karhadkar et al. 2017)

Because the biological functions of sialidases are still unclear, in this report we examined the effects of recombinant human sialidases on human neutrophils. We observed that NEU3 primes neutrophils and changes the localization of CD66, CD11B, CD18, and CD43 in neutrophils. This effect is attenuated in the presence of a NEU3 inhibitor, indicating that this effect is due to NEU3 activity.

## Materials and Methods

### Neutrophil treatment with sialidases and 2AP inhibition assays

Human venous blood was collected from healthy donors who signed written consent, and the procedure was approved by the Texas A&M University Institutional Review Board. Neutrophils were isolated and prepared as previously described (Herlihy et al. 2013; Pilling et al. 2019). 96-well polystyrene plates (353072; BD Biosciences, Franklin Lakes, NJ) were coated with 10 μg/ml human plasma fibronectin (354008; Corning, Corning, NY) in PBS (#17-516F, Lonza, Walkersville, MD) for 1 h at 37°C in a humidified incubator with 5% CO_2_. The plate was then washed three times with PBS (Lonza). 1 μl from a 200 ng/μl stock of NEU1 (TP300386; Origene), NEU2 (TP319858; Origene), NEU3 (Origene), and NEU4 (TP303948; Origene) were each added to 0.1 ml of RPMI-1640 (Corning) to make a 2000 ng/ml solution of each. Serial dilutions of concentrations ranging from 2000 ng to 0.00002 ng for each sialidase were prepared and 100 μl of each sialidase concentration was added to each human fibronectin-coated well. Where indicated, NEU3 was incubated with or without 20 nM or 200 nM of 2AP (A302917, AmBeed, Arlington Heights, IL) for 30 minutes at 37°C in a humidified incubator with 5% CO_2_. 100 μl of 2 × 10^5^ neutrophils in RPMI-1640 (Corning) with 2% BSA (#BSA-50, Rockland, Limerick, PA) was added to each well and left to incubate for 40 minutes at 37°C in a humidified incubator with 5% CO_2_. Differential Interference Contrast (DIC) images of live cells were taken with a 40x objective on a Nikon Ti2 Eclipse microscope. Average circularity of neutrophils was measured as described in (Shihan et al. 2021) using FIJI image J software (Schindelin et al. 2012). For each individual experiment, at least 40 cells were analyzed.

### Sialidase-induced Interleukin-6 accumulation assays

IL-6 accumulation assays were performed as described in (Pilling et al. 2019; Karhadkar, Chen, and Gomer 2020) with the exception that IL-6 levels were measured at 48 hours.

### Activation maker localization assays

Neutrophils were incubated with sialidases as described above for 40 minutes at 37°C in a humidified incubator with 5% CO_2_. 100 µl of the medium was gently removed, and 200 µl of 5% glutaraldehyde (#G-5882, SIGMA, St. Louis, MO) in 1xPBS was added to each well. The cells were fixed and stained as described in (Rijal et al. 2019) using a 1:500 dilution of anti-CD66 (#MAB2244, R&D Systems, Minneapolis, MN), anti-CD62-L (#555542, BD Biosciences), anti-CD11B (#301402, BioLegend, San Diego, CA), anti CD18 (#302102,BioLegend), anti-CD15 (#301902, BioLegend), anti-CD16b (#302002, BioLegend), anti-CD10 (#MCA1556GA, Serotec, Oxford, UK), or anti-CD43 (#555474, BD Biosciences). Secondary antibodies were 1:1000 Alexa fluor 488 Donkey Anti-Mouse IgG (#715-546-150, Jackson ImmunoResearch, West Grove, PA) and actin was stained with Phalloidin-iFluor 555 Reagent (#ab176756, Abcam, Cambridge, UK). Differential Interference Contrast (DIC) and fluorescence images of the cells were taken with a 40x objective on a Nikon Ti2 Eclipse microscope. Cytosolic and cortical localization of the activation markers were analyzed using FIJI image J software (Schindelin et al. 2012) and 40 randomly chosen cells per experiment were analyzed. Phalloidin staining was quantified as described in (Zonderland, Wieringa, and Moroni 2019) using FIJI image J software (Schindelin et al. 2012) and 40 randomly chosen cells per experiment were analyzed.

### Statistical analysis

Statistical analyses were done using Prism version 8.4.1 (GraphPad, San Diego, CA) for *t* tests and one-way or two-way ANOVA with appropriate posttests. Significance was defined as p < 0.05.

## Results

### Human NEU3 alters human neutrophil morphology

Neutrophils circulating in the blood are in a quiescent state with a round, non-adherent morphology (Miralda, Uriarte, and McLeish 2017). As pro-inflammatory signals are released into the blood by local immune cells, neutrophils are first primed, causing them to increase their cell adhesion and develop a more deformable phenotype so that they can leave the circulation and enter the tissue (Condliffe, Kitchen, and Chilvers 1998; Shah, Thomas, and Mehta 2017; Doerschuk et al. 1993; Ekpenyong et al. 2015). To test if NEU3 induces priming or activation of neutrophils, human neutrophils were incubated with recombinant human NEU3, and the live cells were imaged (Figure 1A). Compared to neutrophils not treated with NEU3 (controls), both male and female neutrophils treated with 50 ng/ml of human NEU3 showed a crenulated phenotype and had a decreased average circularity (Figure 1A, B, C, and D). Both male and female neutrophils treated with NEU3 preincubated with either 20 nM or 200 nM of the NEU3 inhibitor 2AP (Karhadkar, Meek, and Gomer 2021) (to generate final 2 AP concentrations of 10 and 100 nM in the presence of neutrophils) had an appearance similar to controls and average circularity not significantly different from controls (Figure 1A, B, C, and D). These data suggest that human NEU3 alters both male and female neutrophil morphology, and that this effect is inhibited with a NEU3 inhibitor.

**Figure 1.**
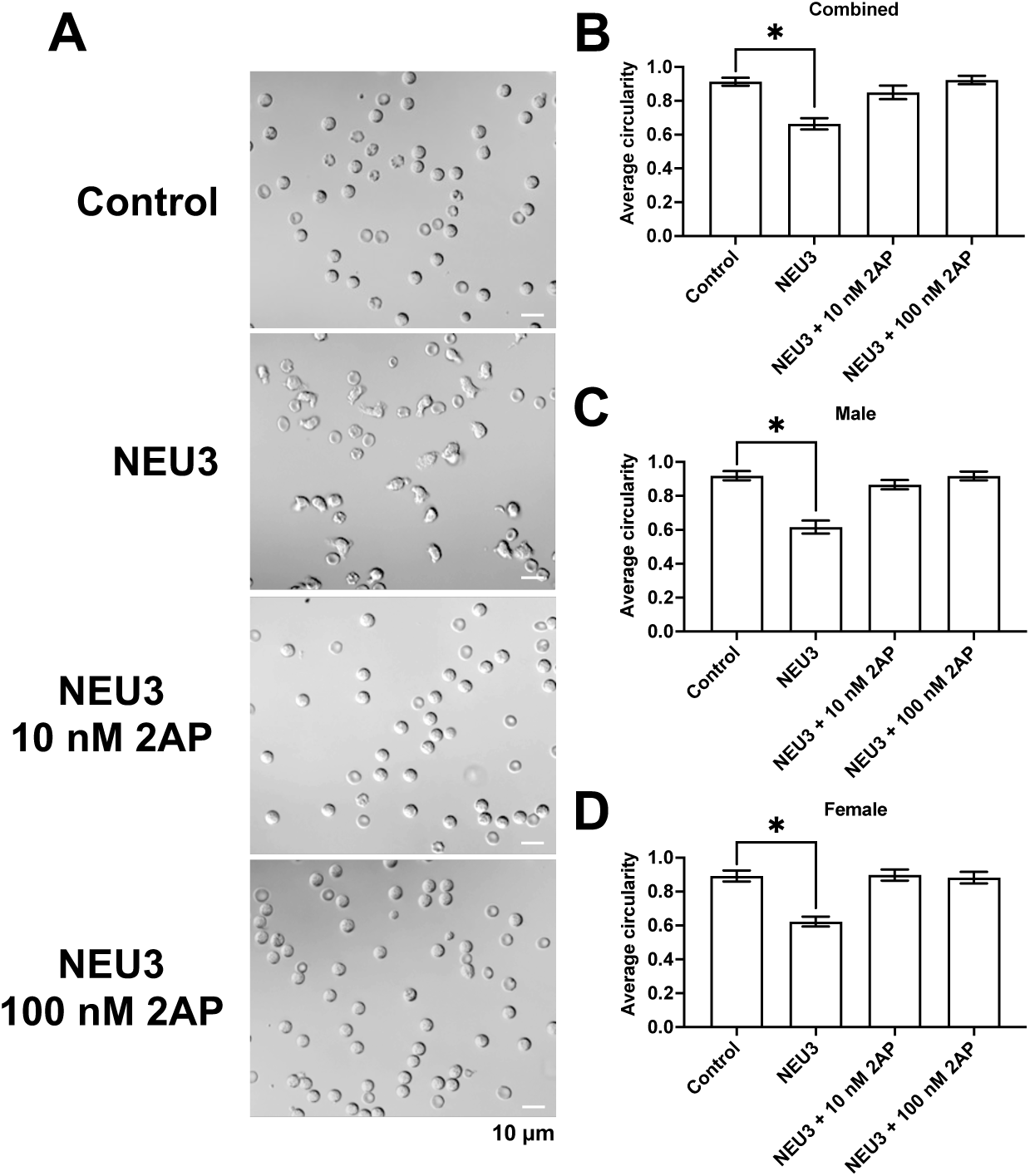
Human NEU3 alters human neutrophil morphology, and NEU3 inhibition attenuates this effect. **A)** Live images of human neutrophils incubated in a uniform concentration of buffer, 50 ng/ml of NEU3, or 50 ng/ml of NEU3 with 100 nM or 10nM of 2AP. DIC images were taken after 40 minutes of incubation. Bar is 10 µm. The smaller toroidal objects are red blood cells. Images are representative of cells from 6 independent experiments (3 male and 3 female donors). Graphs shows average circularity of **B)** combined, **C)** male, or **D)** female neutrophils under the indicated conditions, with at least 50 randomly chosen cells examined for each condition in each experiment. Values are mean ± SEM, n=6. * Indicates p < 0.05 compared to control (unpaired t-tests, Welch’s correction).

### NEU1, NEU2 and NEU4 do not alter neutrophil morphology

To determine if other sialidases affect neutrophil morphology, neutrophils were incubated with sialidases. At concentrations from 0.00001 to 1000 ng/ ml, NEU1, NEU2, NEU4, caused no discernable change in morphology or significant change in average circularity (Figure 2 and Supplementary Figure 1). Neutrophils treated with 100 or 1000 ng/ml of NEU3 showed a crenulated phenotype and decreased average circularity (Figure 2 and Supplementary Figure 1) compared to the control. We also observed that at high concentrations of NEU3, there were no differences in the response of male and female neutrophils (Supplemental Figure 2).

**Figure 2.**
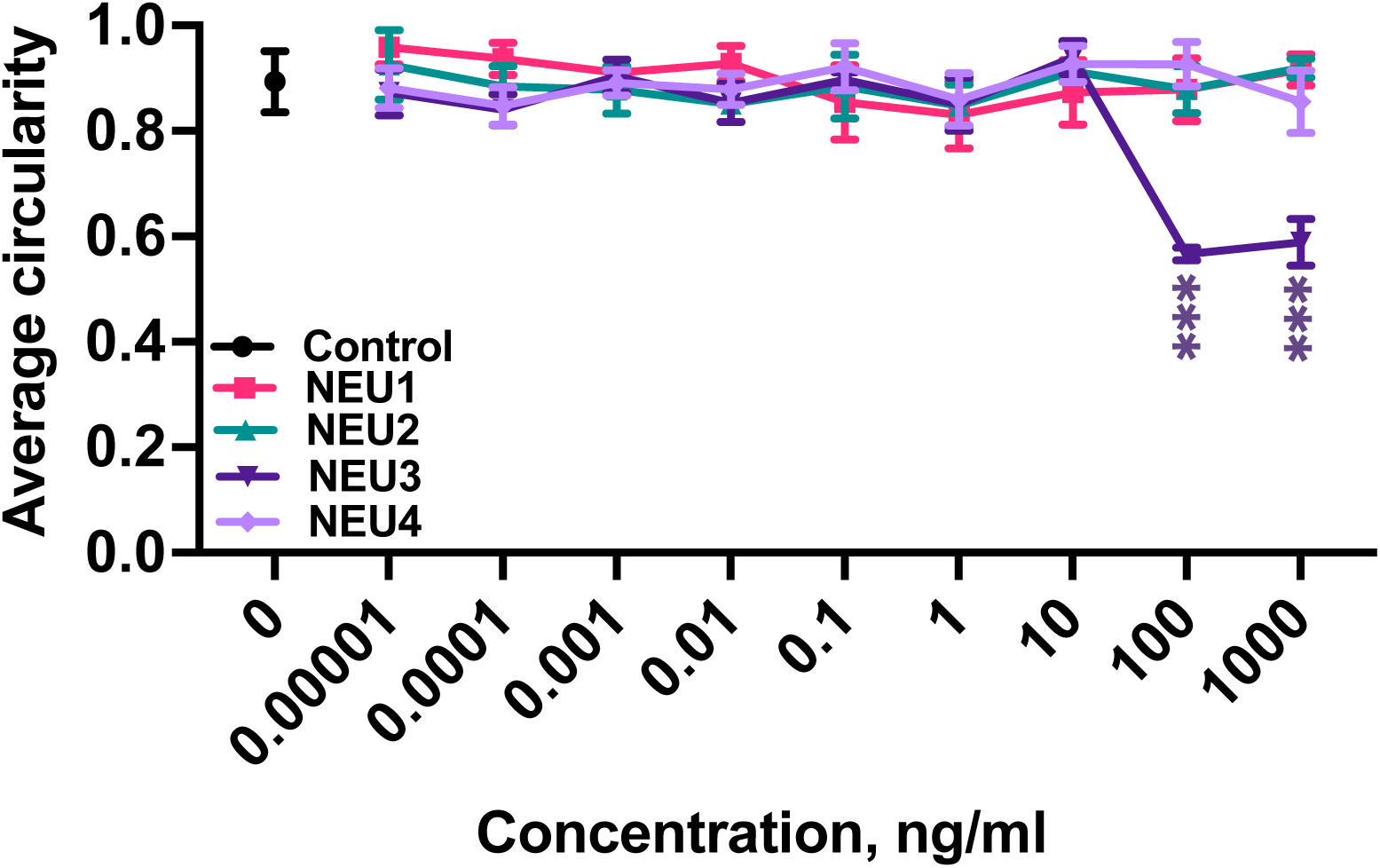
NEU1, NEU2, and NEU4 do not alter neutrophil morphology. Human neutrophils were incubated in different concentrations of the indicated sialidases. Graphs show the average circularity after treatment with NEU1 (magenta), NEU2 (green), NEU3 (dark purple) and NEU4 (light purple). Values are mean ± SEM of the averages from 6 independent experiments (3 male and 3 female donors), with at least 50 randomly chosen cells examined for each condition in each experiment. *** indicates p < 0.001 comparing control to sialidases for each concentration (Unpaired t-tests, Welch’s correction).

To test the hypothesis that the inability of NEU1, 2, and 4 to alter neutrophil morphology was due to these recombinant proteins lacking sialidase activity, we examined the ability of the sialidases to increase the extracellular accumulation of IL-6 by human peripheral blood mononuclear cells (PBMCs) (Karhadkar et al. 2017). As previously observed, NEU1, NEU2, NEU3 and NEU4 increased IL-6 extracellular accumulation (Supplementary Figure 3). This indicates NEU1, NEU2, NEU3 and NEU4 have enzymatic activity, and thus that some specific property of NEU3 in addition to its sialidase activity alters neutrophil morphology.

### NEU3 changes the localization of neutrophil activation markers

Neutrophil priming is a rapid process that balances neutrophil activity and allows for the modulation of response at the inflammation site (Yao et al. 2015; Kitchen et al. 1996). Primed neutrophils undergo both phenotypic and functional alterations (Miralda, Uriarte, and McLeish 2017). Primed neutrophils begin shedding surface selectins to alter their adhesion receptor pattern, undergoing inside-out signaling by promoting exocytosis, leading to the expression of adherence markers such as CD11B, CD35, and CD66B (Yao et al. 2015; Ward, Nakamura, and McLeish 2000; Borregaard et al. 1994), which allow neutrophils to adhere to endothelial cells (Miralda, Uriarte, and McLeish 2017). To determine if NEU1, NEU2, NEU3, or NEU4 induce the expression of activation markers on neutrophils, neutrophils were incubated in buffer or 100 ng/ml of NEU1, NEU2, NEU3, and NEU4. Neutrophils treated with NEU3 had increased expression of CD66, CD11B, and CD18 at the cell cortex and decreased expression in the cytosol (Figure 3 and Supplemental Figure 4). NEU3 treated neutrophils also decreased localization of CD62-L and CD43 at the cortex without a corresponding increase in the cytosol compared to the control (Figure 3 and Supplemental Figure 4). Neutrophils treated with NEU1, NEU2, or NEU4 showed no discernable changes in these markers. Compared to untreated controls, neutrophils treated with all four sialidases showed no discernable changes in the localization of CD10, CD15, and CD16b (Figure 3 and Supplemental Figure 4). Compared to control, neutrophils treated with NEU3 had increased levels of F-actin (Figure 3I and Supplemental Figure 4). NEU1, NEU2, and NEU3 did not increase F-actin levels in the neutrophils (Figure 3I and Supplemental Figure 4). For all conditions, we observed no difference in the localization of the various markers in male and female neutrophils. These data suggest that NEU1, NEU2 and NEU4 do not regulate the localization of activation markers in neutrophils, but NEU3 changes the localization of specific activation markers that may play a role in neutrophil adhesion and migration at inflammatory sites.

**Figure 3.**
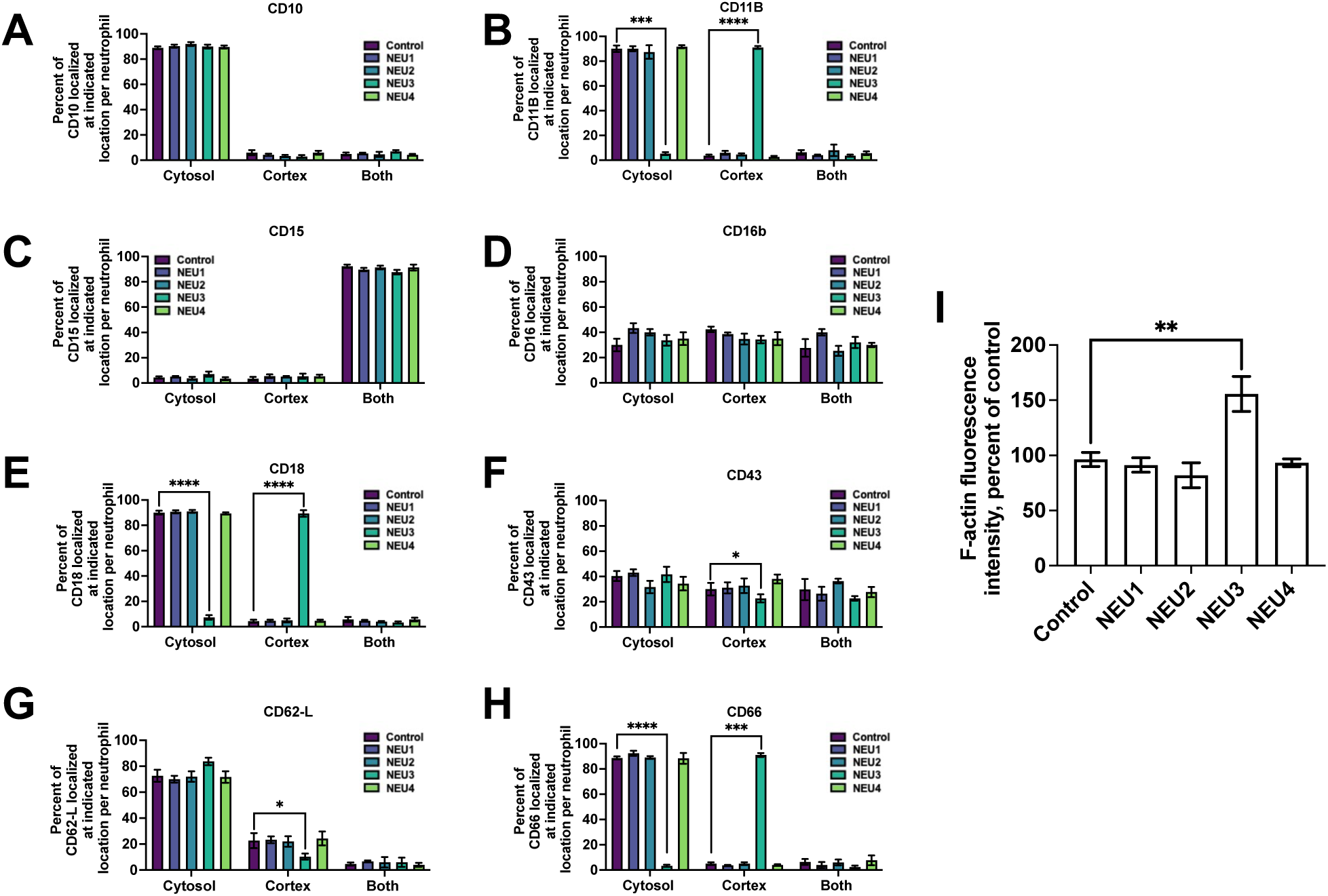
NEU3 changes the localization of some human neutrophil activation markers. Human neutrophils were incubated in the absence (control) or 50 ng/ml of the indicated sialidases. Graphs show the localization of (**A**) C10, (**B**) CD11B, (**C**) CD15, (**D**) CD16b, (**E**) CD18, (**F**) CD43, (**G**) CD62-L, or (**H**) CD66 in either the cytosol, cortex or both in neutrophils. (**I**) Graph shows phalloidin fluorescence staining (a marker for F-actin) intensity in neutrophils after treatment with the indicated sialidases.40 cells were analyzed for each individual experiment. Values are mean ± SEM of 6 independent experiments (3 male and 3 female donors). * indicates p < 0.05 ** p < 0.01 *** p < 0.001, **** p < 0.0001 compared to control (Unpaired t-tests, Welch’s correction).

## Discussion

In this report, we observed that human NEU3 induces phenotypic changes, such as reduced circularity, associated with neutrophil priming and that inhibition of human NEU3 with 10 nM and 100 nM of 2AP attenuates this effect. The total tidal volume of a mouse lung is approximately 0.18 – 0.2 ml (Levitt and Mitzner 1988; Mitzner, Brown, and Lee 2001; Tankersley et al. 1997; Fairchild 1972). We have previously found that at day 21, bronchoalveolar lavage of lungs from bleomycin-treated mice indicated that the lung fluid contained a total of ∽ 15 ng of NEU3, compared to ∼1.7 ng in control mice (Karhadkar, Chen, and Gomer 2020). Assuming that the lung fluid is 10% of the lung volume, this would indicate that in a control mouse lung the NEU3 concentration in the lung fluid is ∼1.7 ng/ (0.1 × 0.2 ml) = ∼85 ng/ ml, while in a fibrotic bleomycin-treated and thus fibrotic lung the NEU3 concentration would be ∼ 750 ng/ ml. We observed that neutrophil priming is induced by treatment of neutrophils with 50 - 1000 ng/ml human NEU3. Further assuming that the concentrations of NEU3 in human lung fluid are similar to the concentrations in mouse lung fluid, this indicates that physiological concentrations of NEU3 in lung fluid induce neutrophil priming. Human NEU1, NEU2, and NEU4 do not appear to prime neutrophils or cause any phenotypic changes associated with neutrophil priming. Human neutrophils incubated with NEU3 increased in CD66, CD11B and CD18 localization at the cortex of neutrophils and decreased in the cytosol. Levels of CD62-L and CD43 were decreased at the cortex but did not have an associated increase in the cytosol of the neutrophils.

Neu3 is upregulated in colon and renal cancer cells (Miyagi et al. 2012), cardiac fibrosis (Ghiroldi et al. 2020), and idiopathic pulmonary fibrosis (Lambre et al. 1988). We have previously detected the upregulation of NEU2 and NEU3 in fibrotic lesions in human and mouse lungs (Karhadkar et al. 2017) and when compared to BAL from control mice, only NEU3 was detected in the BAL fluid from mice with bleomycin-induced pulmonary fibrosis (Karhadkar et al. 2017). The addition of NEU2 and NEU3 to human cells upregulated TGF-β1, and in positive feedback loop, TGF-β1 in turn upregulated the NEU3 expression (Karhadkar et al. 2017; Chen, Lamb, and Gomer 2020). In a recent study, Neu3-null mice had decreased mRNA levels of multiple of immune regulatory chemokines and proinflammatory signals, including interleukin-1β, TNF-α, and TGF-β (Yang et al. 2021), coupled with a significantly reduced leukocyte infiltration, including neutrophils (Yang et al. 2021). Together with our findings, this reveals that human NEU3 might be a regulator of neutrophil priming by directly or indirectly influencing the accumulation and activation of neutrophils at a site of inflammation.

Compared to quiescent circulating neutrophils, neutrophils experience both morphological changes and molecular changes in the presence of priming agents. Primed neutrophils also have an increased expression of adhesive markers on the membrane and have increased intracellular reactive oxidative species production (Ekpenyong et al. 2017). Priming enhances neutrophil phagocytosis and degranulation (Kitchen et al. 1996; Vadas et al. 1984; Cross and Wright 1991). Primed neutrophils from patients with diabetes have been found to be more rigid than those from healthy patients (Pecsvarady et al. 1994). A study of anti-inflammatory drugs used neutrophil deformability, morphological changes, and increased neutrophil elastase as measures of neutrophil activation (Craciun et al. 2013).

Adhesion of primed neutrophils to the endothelial lining is important to induce outside-in signaling (Nair and Zingde 2001), allowing for the expression of adhesive markers on the surface of neutrophils (Vogt et al. 2018; Condliffe, Kitchen, and Chilvers 1998). The CD66 are glycoproteins that play a role in neutrophils adhesion to E-selectin on local endothelial cells through the presentation of CD15s epitope (Kuijpers, Hoogerwerf, van der Laan, et al. 1992). CD66 expression on neutrophils also promotes neutrophil adhesion to fibronectin through the expression of β2 integrin (Stocks et al. 1996). When neutrophils undergo priming after exposure to fMLP, CD66 expression initiates an increase in superoxidase production (Stocks et al. 1996; Ruchaud-Sparagano et al. 1997). We observed that when neutrophils are treated with 50 ng/ml of human NEU3, CD66 localization increases at the cortex of the neutrophils indicating the initiation of neutrophil adherence and priming. CD62-L is known as the tethering or rolling receptor (Ivetic, Hoskins Green, and Hart 2019). Neutrophils that are primed or activated shed CD62-L and in turn, increase CD11B (Simon et al. 1995). Neutrophils at infection sites had an upregulation in CD11B and reduced expression of CD62-L (Kuhns, Long Priel, and Gallin 1995; Choi et al. 2003). Patients with diabetic microangiopathy had decreased surface expression of CD62-L on neutrophils (Mastej and Adamiec 2008). We observed that when neutrophils are treated with 50 ng/ml of human NEU3, CD62-L expression at the cortex decreases with no increase in the cytosol indicating that upon neutrophil priming, CD62-L begins to shed. CD11B and CD18 are primarily expressed on a variety of immune cells, including neutrophils (Weirich et al. 1998; Kim et al. 2003; Abdel-Salam and Ebaid 2014). On neutrophils, CD11B and CD18 are noncovalently bound to form a functional heterodimer (Khan, Khan, and Gupta 2018). The cortical localization of this heterodimer is involved in regulating multiple leukocyte responses such as adhesion and migration (Plow and Zhang 1997). We observed that on neutrophils treated with 50 ng/ml of human NEU3, CD11B and CD18 had increased localization at the cortex and decreased localization in the cytosol, indicating that NEU3 promotes neutrophil adhesion. In patients with acute or chronic inflammatory conditions, CD10 is widely utilized to discriminate mature and immature neutrophil populations (Marini et al. 2017; Elghetany 2002; Cossman et al. 1983; McCormack, Nelson, and LeBien 1986). Although the function of CD15 is not well known, it has been observed to be expressed on mature neutrophils and is involved in multiple neutrophil functions such as cell-cell interactions, phagocytosis, and degranulation (Nakayama et al. 2001; Melnick et al. 1986; Melnick et al. 1985; Skubitz and Snook 1987; Forsyth, Simpson, and Levinsky 1989; Warren et al. 1996).

CD16b is specifically expressed on neutrophils (Zhang 2000). CD16b plays a role in neutrophil degranulation (Unkeless et al. 1995), phagocytosis (Fernandes et al. 2005), and calcium mobilization (Wang and Jonsson 2019). We observed that in neutrophils treated with 50ng/ml of human NEU3, there was no change in the localization of CD10, CD15, or CD16b, indicating that these activation markers might be regulated at a later stage in neutrophil priming or activation. CD43 is found specifically on neutrophils and in very rare cases, on colon carcinomas and on several non-hematopoietic cell lines (Wong et al. 1990; Fernandez-Rodriguez et al. 2002). Similar to CD62-L, CD43 is shed from the membrane upon neutrophil activation or by TNF-α priming (Campanero et al. 1991; Kuijpers, Hoogerwerf, Kuijpers, et al. 1992; Rieu et al. 1992; Remold-O’Donnell and Parent 1994) during adhesion and spreading (Nathan et al. 1993; Lopez et al. 1998). When neutrophils were treated with 50 ng/ml of human NEU3, CD43 localization at the cortex decreased with no increase in the cytosol, suggesting that NEU3 induces shedding of CD43 on neutrophils.

The broad range of neutrophil responsiveness confers extensive functional flexibility, allowing neutrophils to respond rapidly and appropriately to varied and evolving threats throughout the body. Uncontrolled priming and activation of neutrophils may lead to multiple inflammatory disorders, such as COPD, arthritis, and IPF. The ability of NEU3 to induce neutrophil priming suggests that inhibiting NEU3 activity, or NEU3 upregulation, may be beneficial to patients with a variety of diseases.

## Figure Legends

**Supplementary Figure 1.**
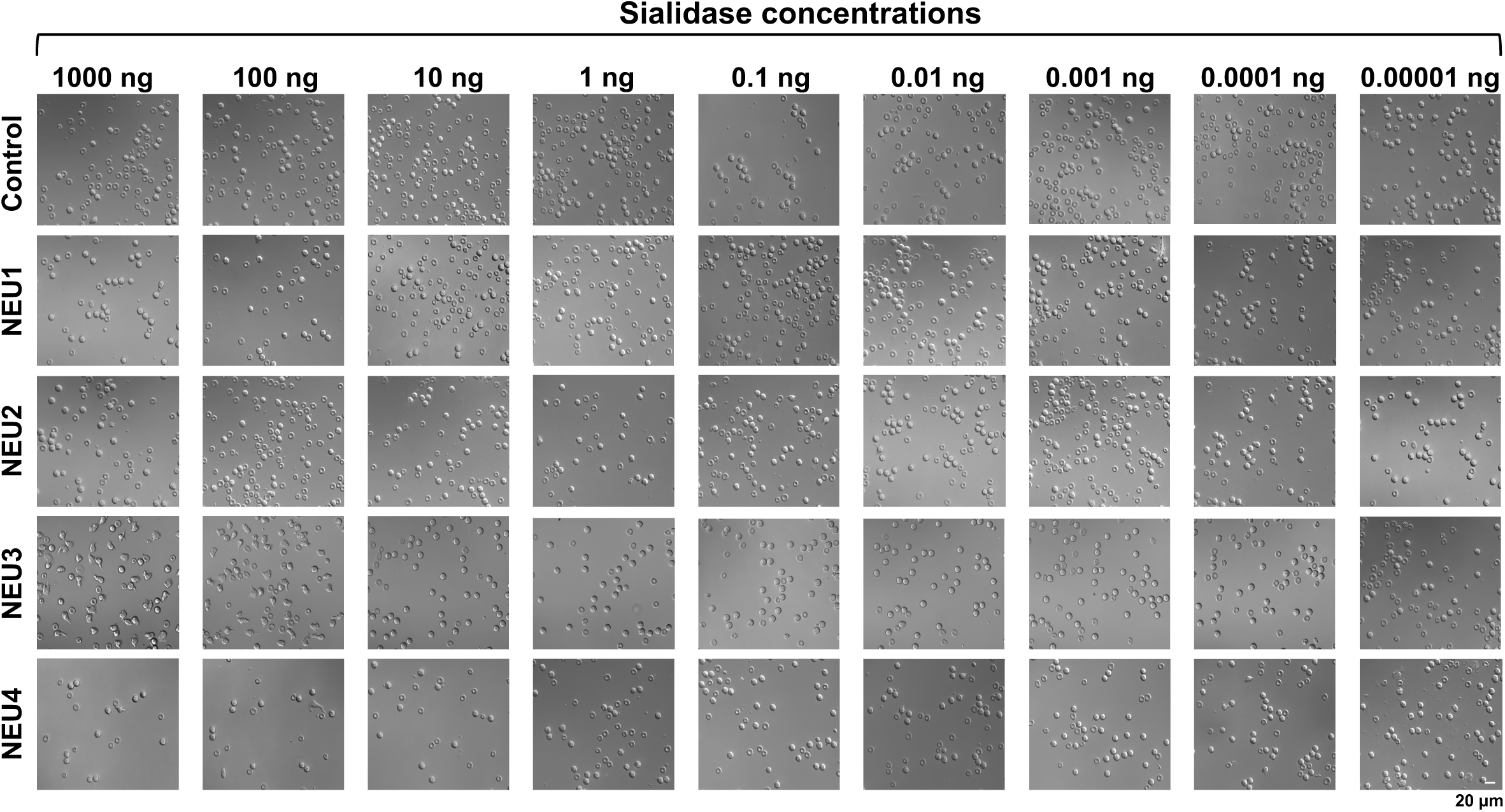
NEU1, NEU2, and NEU4 do not discernably alter neutrophil morphology. DIC images show human neutrophils incubated in different concentrations of the indicated sialidases. After 40 minutes, cells were imaged. Bar is 20 µm. The smaller toroidal objects are red blood cells. Images are representative of 6 individual experiments.

**Supplementary Figure 2.**
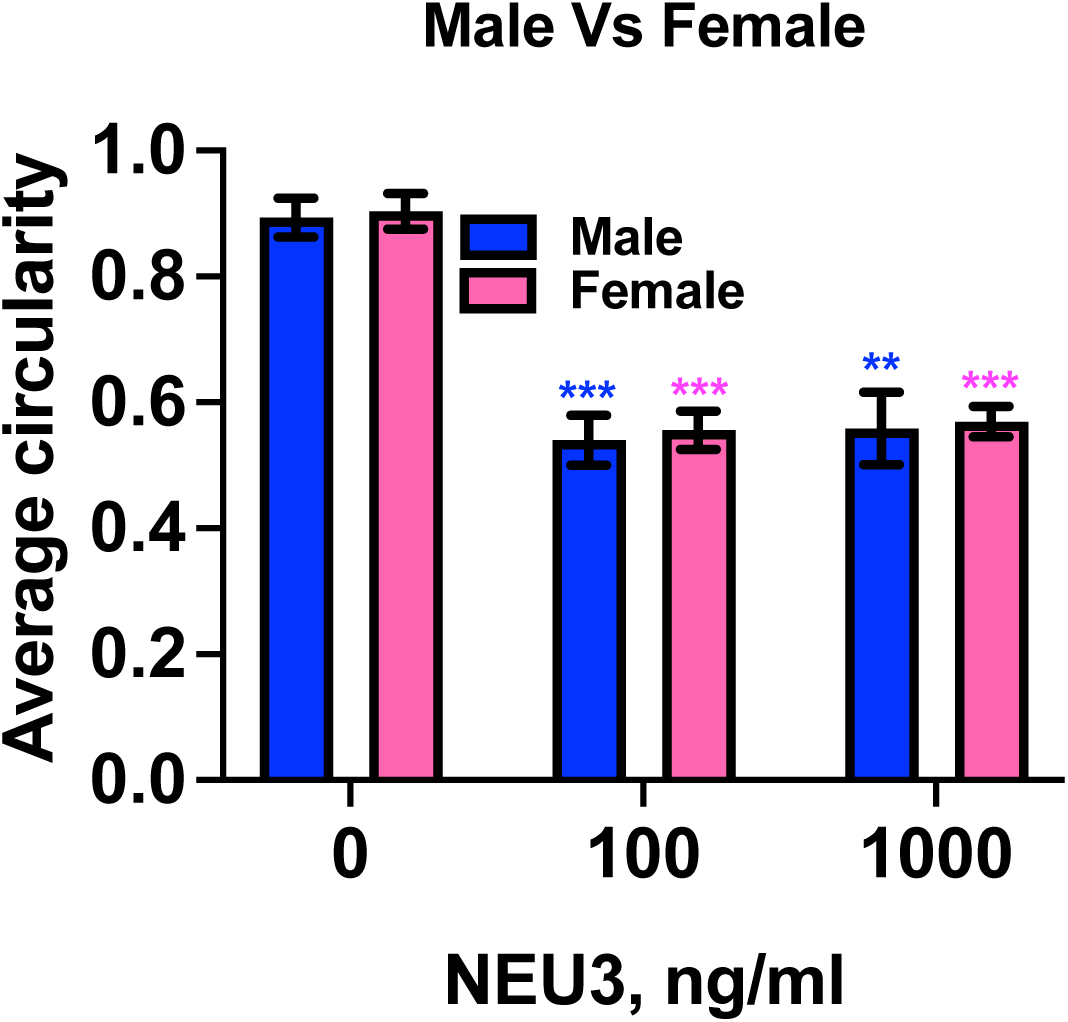
Both male and female human neutrophils show a decrease in circularity after NEU3 treatment. Human male and female neutrophils were incubated with the indicated concentrations of NEU3. Graphs show the average circularity of male (blue) and female (pink) neutrophils after treatment with NEU3. Values are mean ± SEM of the averages from 6 independent experiments (3 male and 3 female donors), with at least 50 randomly chosen cells examined for each condition in each experiment. ** indicates p < 0.01, *** indicates p < 0.001 compared to 0 NEU3 control (2-way ANOVA, Holm-Šídák’s test).

**Supplementary Figure 3.**
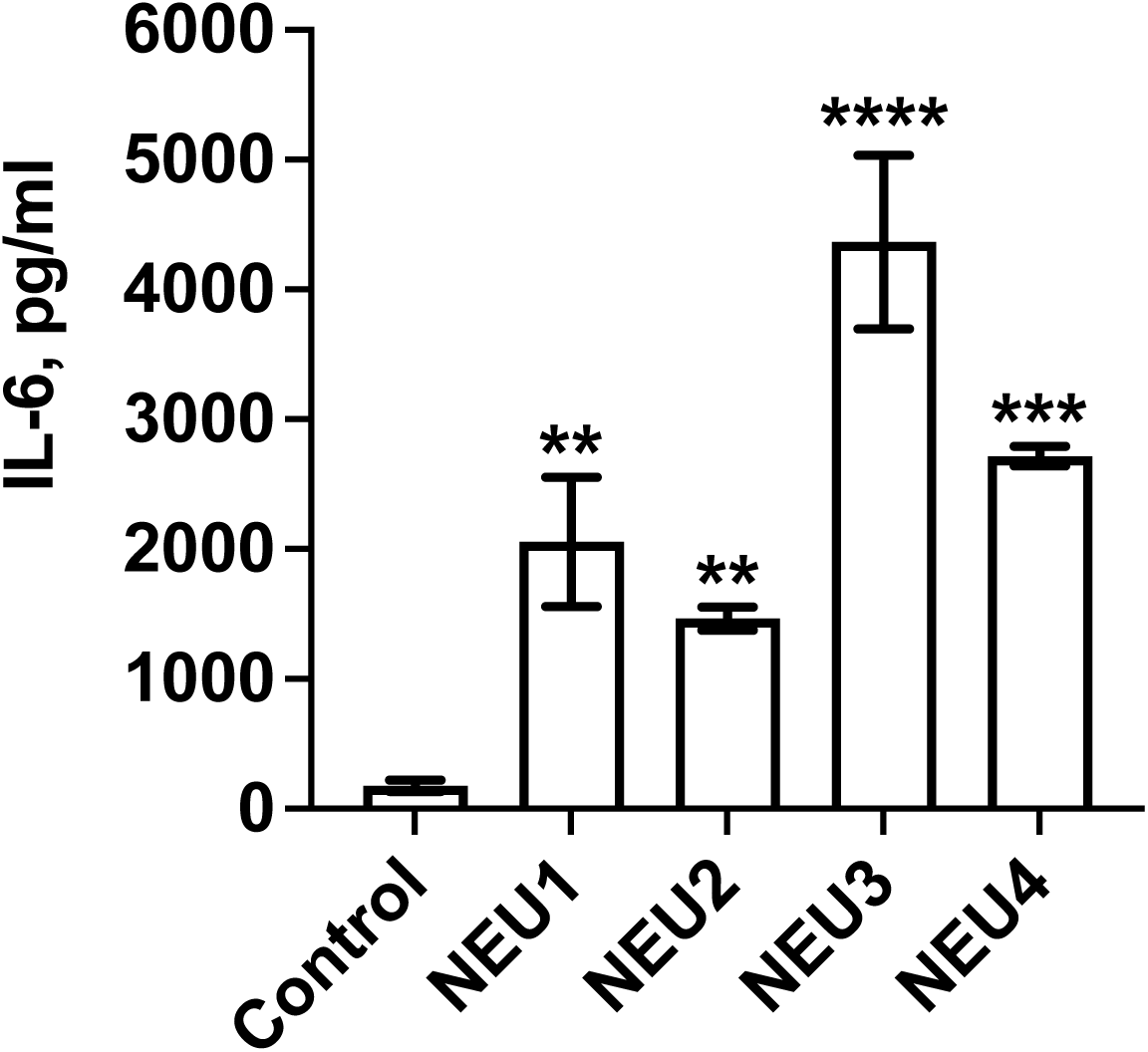
The four sialidases have bioactivity. Human PBMCs were incubated with the indicated recombinant human sialidases for 48 hours. Supernatants were assayed for IL-6 by ELISA. Graph shows amount of accumulated extracellular IL-6. Values are mean ± SEM of 2 independent experiments. ** indicates p < 0.01 *** p < 0.001, **** p < 0.0001 compared to no sialidase control (unpaired t-tests, Welch’s correction).

**Supplementary Figure 4.**
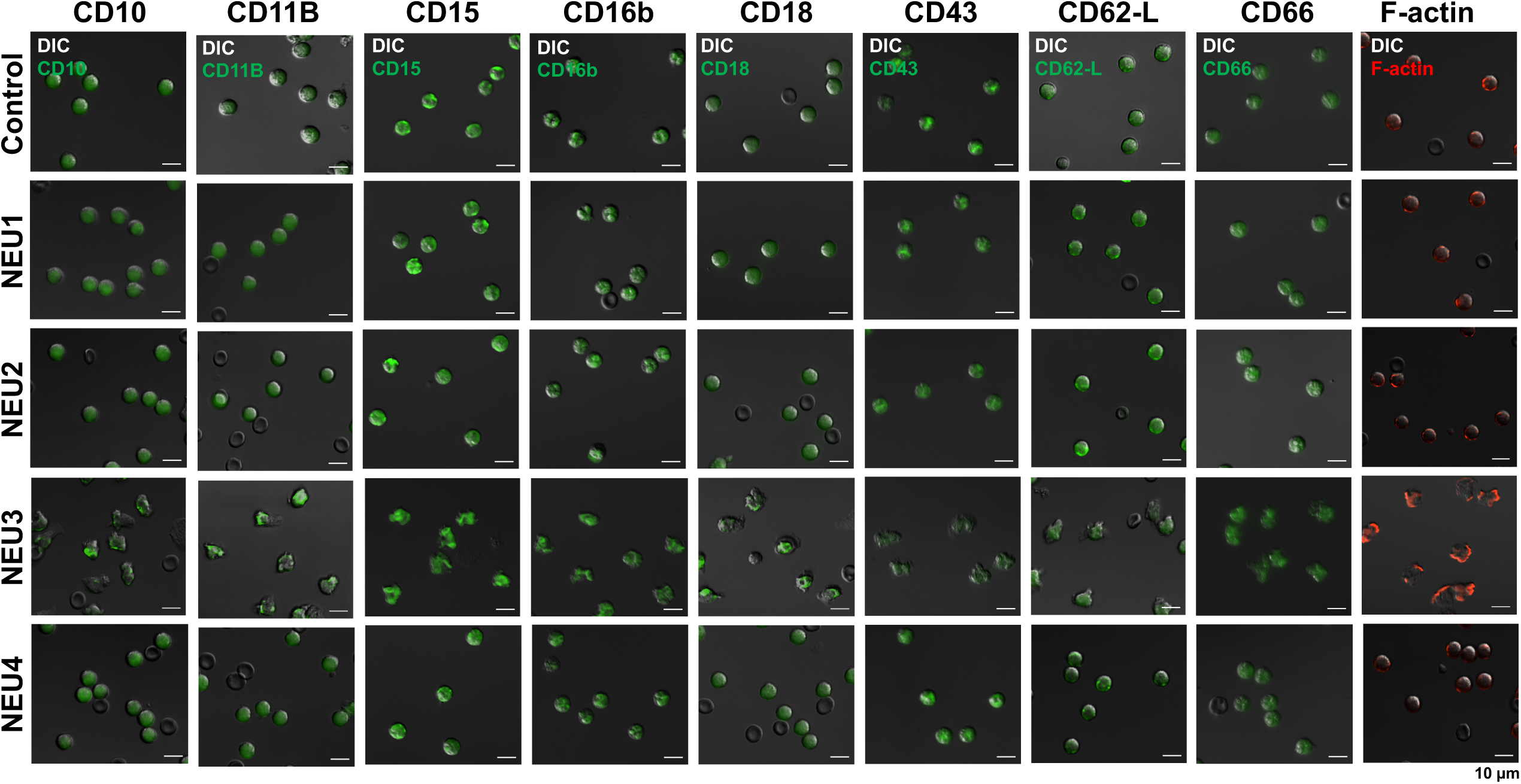
Localization of markers after treatment with sialidases. Human neutrophils were incubated in the absence (control) or presence of 50 ng/ml of the indicated sialidases. After 40 minutes, cells were fixed and stained. DIC and fluorescent images were taken with a 40x objective. Bar 10 µm. Images are representative of 6 individual experiments (3 male and 3 female donors).

